# Synergy with TGFß ligands switches WNT pathway dynamics from transient to sustained during human pluripotent cell differentiation

**DOI:** 10.1101/406306

**Authors:** Joseph Masseya, Yida Liua, Omar Alvarengaa, Teresa Saeza, Matthew Schmerera, Aryeh Warmflasha

**Author notes:** present address: Centers for Disease Control, Atlanta, Georgia, 30333 (Matthew Schmerera). present address: Sainsbury Laboratory, Cambridge University, UK.

## Abstract

WNT/ß-catenin signaling is crucial to all stages of life. It controls early morphogenetic events in embryos, maintains stem-cell niches in adults, and is disregulated in many types of cancer. Despite its ubiquity, little is known about the dynamics of signal transduction or whether it varies across contexts. Here we probe the dynamics of signaling by monitoring nuclear accumulation of ß-catenin, the primary transducer of canonical WNT signals, using quantitative live-cell imaging. We show that ß-catenin signaling responds adaptively to constant WNT signaling in pluripotent stem cells, and that these dynamics become sustained upon differentiation. Varying dynamics were also observed in the response to WNT in commonly used mammalian cell-lines. Signal attenuation in pluripotent cells is controlled by both intra- and extra-cellular negative regulation of WNT signaling. TGFß-superfamily ligands Activin and BMP, which coordinate with WNT signaling to pattern the gastrula, increase the ß-catenin response in a manner independent of their ability to induce new WNT-ligand production. Our results reveal how variables external to the pathway, including differentiation status and crosstalk with other pathways, dramatically alter WNT/ß-catenin dynamics.

## Introduction

During metazoan development, WNT/ß-catenin signaling is critical for proper morphogenesis and patterning of tissues and cell-types (1, 2). In the adult, WNT plays a key role in maintaining homeostasis and regulating stem cell niches (3, 4). Mutations in WNT/ß-catenin pathway components are frequently found in human cancers (5, 6).

Canonical WNT signals are transduced to the nucleus by ß-catenin (**Fig. 1*A***) (6–8). WNTs are secreted lipid-modified proteins which bind a cell-surface receptor complex consisting of Frizzled and LRP5/6 (7, 9, 10). WNT-receptor binding leads to inhibition of the ß-catenin destruction complex (consisting of multiple scaffolds and kinases, including APC, AXIN, GSK3ß and CK1) (11), allowing ß-catenin accumulation and translocation to the nucleus. In the nucleus, ß-catenin complexes with TCF/LEF to regulate target genes in a context-specific manner (12). In addition to being the primary effector of canonical WNT signaling, ß-catenin also has a role stabilizing adherens junctions at the cell membrane.

**Figure 1.**
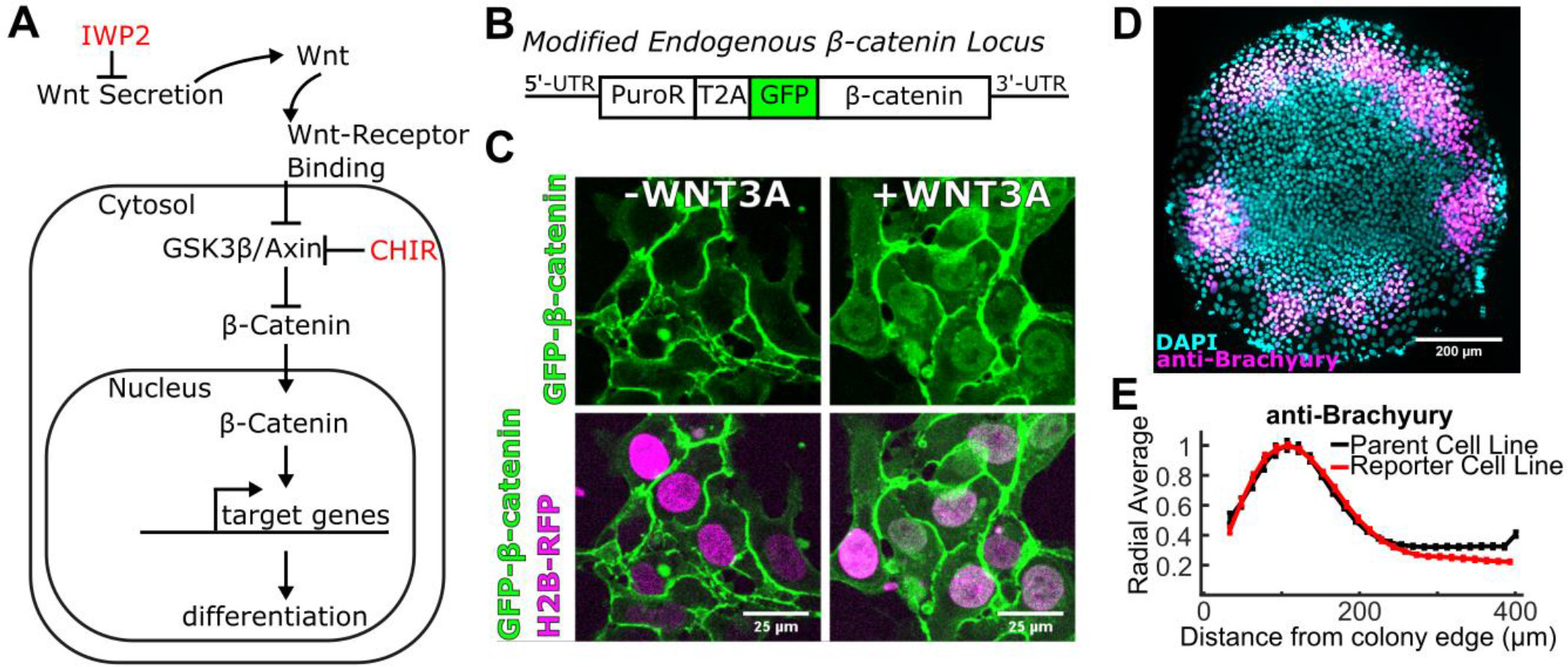
A CRISPR-Cas mediated GFP-knock-in labels endogenous β-catenin and does not disturb signaling or differentiation. (A) Simplified canonical WNT/β-catenin pathway. Pharmacological small molecules used in this study are shown in red. (B) Schematic of resulting mRNA of labeled ß-catenin allele after CRISPR-cas mediated PuroR-T2A-GFP knock-in. The puromycin resistance protein (PuroR) facilitates selection of labeled cells. T2A is a self-cleaving peptide which allows PuroR to be separated from GFP-ß-catenin. (C) Confocal microscopy images of live hESCs with GFP-labeled β-catenin and the nuclear marker H2B-RFP. Left image is prior to exogenous WNT3A addition. Right image shows nuclear GFP-β-catenin accumulation after 4 hours of WNT3A treatment. (D) A representative image of an 800 micron diameter micropatterned hESC colony stained with DAPI (magenta) and Brachyury (cyan) using immunocytochemistry after 42 hours of treatment with exogenous BMP4. (E) Quantification of BRA expression in the hESC cell-line and the labeled reporter cell-line, respectively (N = 20 for unlabeled cells and N=11 for reporter cells).

During gastrulation, dramatic morphogenetic rearrangements occur simultaneously with the patterning of the primitive streak by BMP, WNT, and NODAL signals (13). Given these cellular movements and the rapid changes in expression patterns of all of these ligands, it is clear that cells will experience rapidly changing levels of all of these morphogens. The coupling between patterning, growth, and morphogenesis, along with the lack of methods for temporally precise perturbation of signaling, makes systematically dissecting the contribution of signaling dynamics difficult *in vivo*. In contrast, *in vitro*, researchers can administer precise amounts of signaling ligands while inhibiting endogenous ligands. Similarly, the combinatorial effects of multiple ligands can be directly investigated. Finally, ligands can be provided dynamically, which allows for testing effects of different ligand dynamics such as adding the same doses of ligand at different rates-of-change (14, 15). Additionally, *in vitro* cell-culture is highly amenable to live-cell imaging techniques.

While numerous regulators of the WNT/ß-catenin pathway have been identified (7, 16, 17), less is known about WNT/ß-catenin signaling dynamics. Because of the variety of contexts in which it plays crucial roles, and the diversity of potential regulators, it is impossible to understand ß-catenin dynamics in any particular setting without making explicit measurements. Here we created a fusion of GFP and ß-catenin at the endogenous locus and utilized quantitative microscopy to measure signaling dynamics. We found that the response to WNT varies significantly by differentiation stage and cell-type. ß-catenin response was adaptive to WNT in human embryonic stem cells, but sustained in many other cell-types. Adaptation in hESCs is controlled upstream of GSK3 by both intra- and extra-cellular mechanisms, and confers sensitivity to the WNT rate of change at lower doses. However, when hESCs were subjected to a primitive-streak differentiation protocol (18), ß-catenin was stably activated. Surprisingly, TGFß and BMP both synergized with exogenously provided WNT by a mechanism independent of WNT-ligand induction, and BMP could induce nuclear ß-catenin independently of WNT ligands altogether. Our results reveal new insights into the how WNT/ß-catenin signaling dynamics vary by context, and how WNT signaling synergizes with other key morphogens during early development.

## Results

### A CRISPR-cas mediated GFP-knock-in labels endogenous β-catenin without perturbing signal transduction or differentiation

In order to measure WNT/ß-catenin signaling dynamics in single cells, we used CRISPR-Cas9 gene editing (19–22) to insert GFP at the N-terminus of endogenous ß-catenin in hESCs (**Fig. 1*B***,***C***). To facilitate computational nuclear identification and analysis, these cells additionally express RFP-H2B incorporated into the genome with the ePiggyBac transposable element system (23). GFP-ß-catenin localization was confirmed with immunofluorescent antibody staining of ß-catenin (**Fig. S1*A***), and correct expression of the GFP-ß-catenin fusion without any other expression of GFP was confirmed by Western blotting (**Fig. S1*B***). GFP-ß-catenin accumulated in the nucleus in response to exogenous WNT3A (**Fig. 1*C***), and the transcriptional dynamics of β-catenin target genes in response to WNT3A were highly similar in unmodified and modified hESCs (**Fig. S1*C***). As a stringent test of the potential of these cells, spatial patterning and differentiation were unaffected in a micropatterned gastruloid protocol (24, 25) (**Fig. 1*D*,*E***).

### Human pluripotent cells respond adaptively to exogenous WNTs, and adaptation is controlled upstream of GSK3

In hESCs treated with exogenous WNT3A, GFP-ß-catenin initially rapidly accumulates in both the nucleus and cell membrane (**Fig. 2*A***,***B***). However, while the increase in membrane GFP-β-catenin is sustained, the nuclear GFP-β-catenin begins to decline approximately 4 hours after WNT3A addition (i.e., hESCs adapt to constant WNT signals). Peak signaling is dose dependent and occurs earlier at lower doses. Adaptation is complete at sufficiently low doses of WNT3A, and at saturating WNT3A (doses above 300ng/ml) cells adapt to approximately 40% of peak signal. A recently developed, commercially available iPSC line with a GFP-ß-catenin fusion (Allen Institute) had identical dynamics (**Fig. S*2***). Signaling dynamics were verified in unmodified hESCs using immunofluorescent antibody staining for ß-catenin (**Fig. 2*C***).

**Figure 2.**
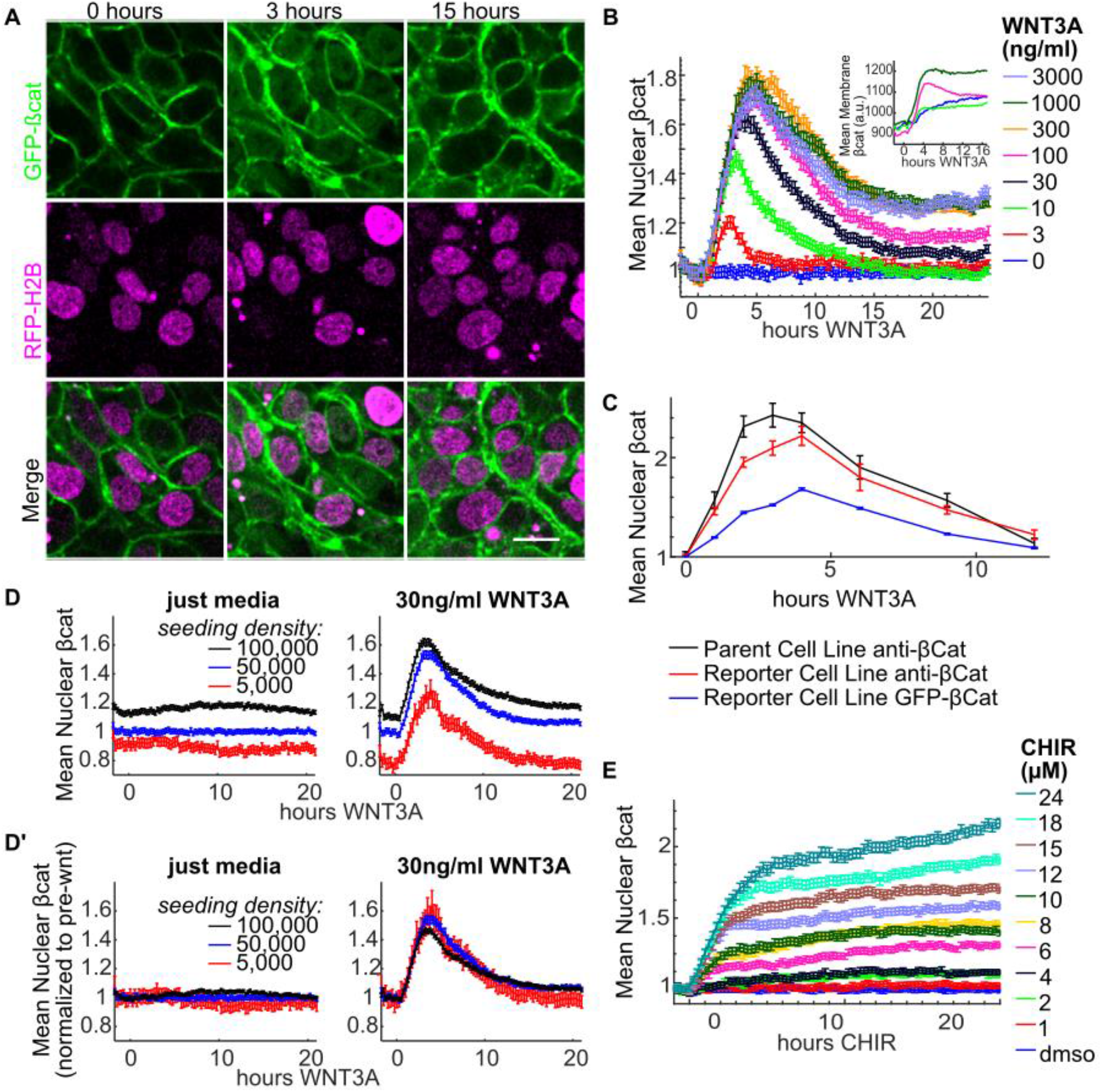
WNT/β-catenin signaling is partially adaptive in stem cells, and adaptation is controlled upstream of GSK3ß. (A) Representative images from time-lapse imaging of GFP-ß-catenin labeled hESCs treated with 100ng/ml WNT3A at 0, 3, and 15 hours. (B) Quantification of nuclear and membrane (insert) levels of GFP-ß-catenin in hESCs treated with varied concentrations (ng/ml) of WNT3A. (C) Quantification of ß-catenin dynamics by either GFP or immunofluorescence in the reporter cell line compared with immunofluorescence in the parental line. (D) GFP-ß-catenin labeled hESCs at indicated seeding densities with or without WNT3A, represented as ratio to 50,000cells/cm_2_. This is seeding density for all other experiments. (Dʼ) Same data as in D, but represented as ratio to mean signaling prior to WNT addition at that density. (E) GF-ß-catenin labeled hESCs treated with varied concentrations (μM) of CHIR99021, a pharmacological GSKβ inhibitor. Error bars in all graphs indicate SEM of ≥ 644 cells. In this and all other figures, nuclei are computationally identified using a nuclear label (RFP-H2B if live-cells, DAPI if not). The mean intensity of the indicated marker (GFP-ß-catenin, anti-ß-catenin) is normalized against the intensity of the nuclear label. Except for D̓, "gMean nuclear βcath" is always defined as the ratio over the negative control for that experiment. Membrane (a.u.) is the mean pixel intensity of computationally identified membranes across at least 8 images per condition after background subtraction. We note that the values shown for nucleus and membrane are not directly comparable as only the former are normalized. Scale bar is 20μm.

Post-translational palmitoleoation increases hydrophobicity of WNT proteins and is thought to make them prone to aggregation and increase difficulty of purification (26, 27). Additionally, recombinant WNT is likely not as potent as its endogenous counterpart, as exogenous doses used *in vitro* are very high (often more than 200ng/ml) when compared to other morphogens (Activin and BMP4 both saturate Smad signaling at 3ng/ml (15)). This has lead other researchers to question the stability of recombinant WNTs in culture media (28–30). However, if the observed adaptation to WNT signals was due to WNT3A instability in culture-media, the time to adapt to WNT should be dose dependent and become very long at doses well above saturation, which is not the case. Additionally, at saturating WNT3A, conditioned media taken from WNT3A-treated hESCs retains its ability to activate signaling in previously unstimulated cells over the entire course of adaptation (**Fig. S3**), proving that adaptation at this dose is not due to WNT instability. Finally, a commercially available “WNT-stabilizer” (31), had no effect on the adapatation of hESCs (**Fig. S4*B***). We note that this WNT-stabilizer caused cells to quickly aggregate (**Fig. S4*A***) and increased their nuclear area (**Fig. S4*C***), which likely indicates biological effects beyond simply stabilizing recombinant WNT.

To assay possible effects of endogenous WNT ligands induced by the exogenously supplied WNT3A, cells were treated with a small molecule inhibitor of WNT secretion, IWP2 (32), along with either a high or intermediate dose of exogenous WNT3A (**Fig. S5**). Inhibition of WNT secretion lowered the final adapted level only at intermediate doses, indicating that additional WNT ligands are induced, but the high dose of WNT3A alone is capable of completely saturating the response, and is not additive with endogenous WNT signaling. Additionally, while increasing cell density increased the total GFP-ß-catenin in cells (**Fig. 2*D***), it did not affect the fold change in nuclear ß-catenin in response to stimulation (**Fig. 2*D’***), nor were dynamics affected by ROCK inhibition with the small molecule Y-27632 (**Fig. S6**). Pathway activation using the small molecule GSK3ß inhibitor, CHIR99021 (commonly used as a WNT substitute in differentiation protocols) (18, 33–35), resulted in non-adaptive dose-dependent increases of nuclear ß-catenin (**Fig. 2*E***). Sustained signaling in response to CHIR99021 demonstrates that the mechanisms that control adaptation must act upstream of GSK3ß. This result also highlights that while small molecule inhibitors of GSK3ß are potent activators of ß-catenin signaling, they induce very different dynamics than WNT ligands.

### WNT adaptation is governed by intra- and extra-cellular mechanisms

We next aimed to separate the effects of potential adaptive mechanisms that act intra-cellularly (e.g., negative feedback at the destruction complex) from those that act extra-cellularly (eg., negative feedback from secreted WNT-inhibitors like DKKs, SFRPs, WIFs, and Notum). We hypothesized that if adaptation involved extra-cellular negative regulators, conditioned media taken from WNT-treated adapting cells would progressively lose its ability to activate previously unstimulated cells. To test this hypothesis, we treated cells with varying doses of WNT3A for 8 hours, by which time they had begun to adapt, and then swapped the media from these cells with that of previously untreated cells while measuring the resulting signaling in each group (**Fig. 3*A***). The response was greatly decreased in the second group of cells when sub-saturating WNT was used. However, at saturating WNT, the response of the second group was essentially unchanged, and this persisted over the entire course of adaptation (**Fig. S3**). Thus, the media undergoes a net decrease in WNT/ß-catenin activating potential, but only when a sub-saturating dose of WNT is used. In a complimentary experiment, we periodically replaced WNT3A-containing media with fresh media containing the same dose of WNT3A (thus replacing inactivated WNT and potentially removing secreted regulators). At subsaturated doses, replenishing the WNT lead to a more sustained response, while cells treated with high WNT were unaffected by media replacement (**Fig. 3*B***). This evidence demonstrates that extra-cellular adaptation mechanisms exist, but are only functional at sub-saturating WNT. Intra-cellular mechanisms must also exist since adaptation at saturating WNT can not be explained by extra-cellular mechanisms.

**Figure 3.**
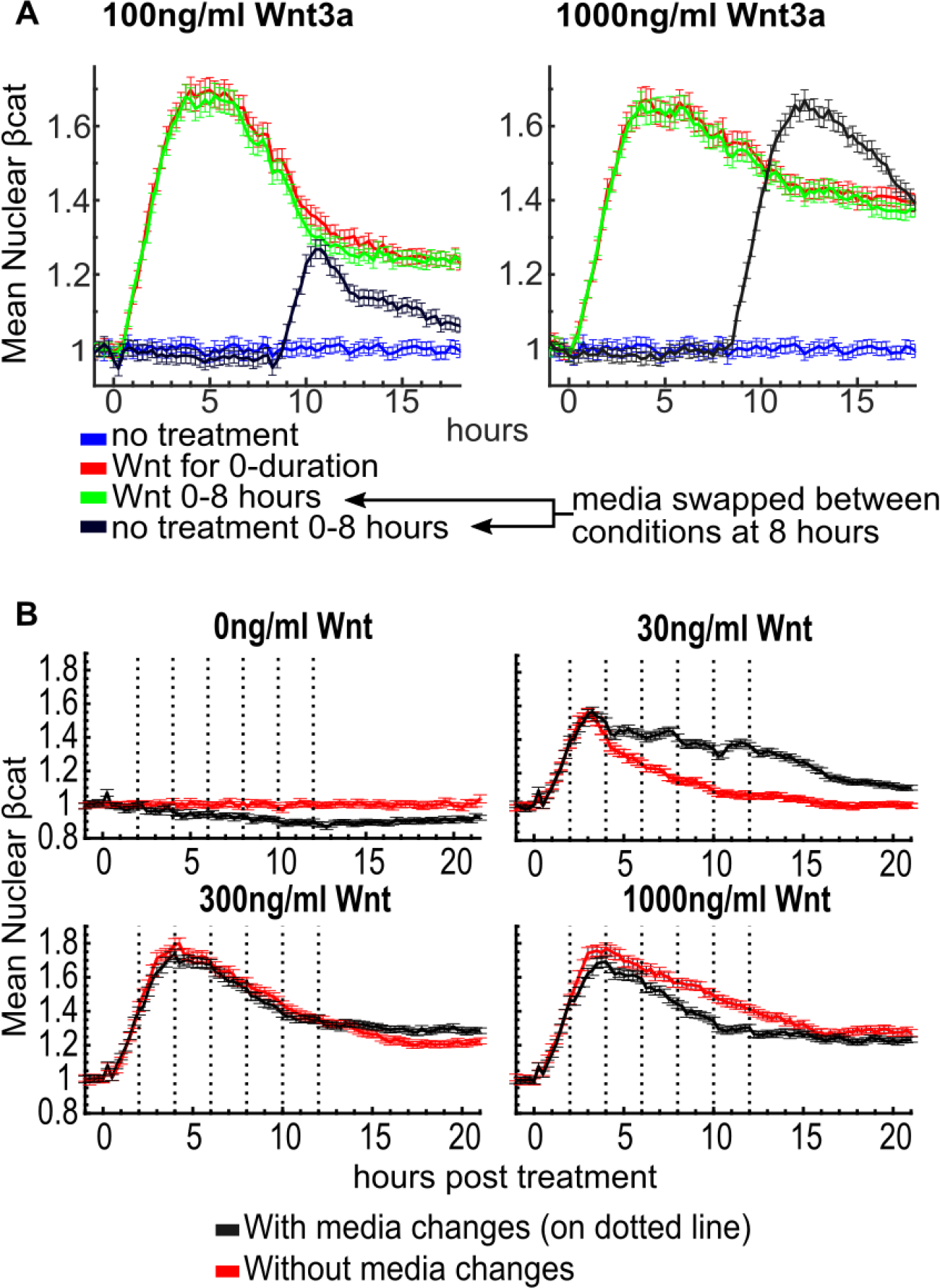
Adaptation mechanisms act both intra- and extra-cellularly. (A) Quantification of ß-catenin dynamics with live cell imaging. At 0h, cells were either left untreated (blue and black curves) or treated with 100ng/ml WNT3A (left) or 1000ng/ml WNT3A (right) (red and green curves). After 8 hours, media was either swapped between a treated (green) and untreated (black) well or not (red and blue). The experiments on the left and right panels were performed simultaneously so that a single no treatment control (blue) was used and is reproduced in each graph. (B) Cells were treated with indicated concentrations of WNT3A. They were then either left unperturbed (red) or had their media replaced with fresh media containing an identical concentration of WNT3A (black) at the indicated time-points (dashed gray line; every 2 hours for initial 12 hours). Error bars in all graphs indicate SEM of ≥ 494 cells.

### hESCs are sensitive to WNT dynamics

Previous work with the TGFβ and Shh pathways has shown that ligand presentation dynamics can alter the signaling response and change cell fate decisions (14, 15, 36–38). Hence, we tested whether hESCs are sensitive to WNT dynamics. A hallmark of signaling adaptation is sensitivity to the rate of ligand presentation (i.e., the time derivative of the ligand concentration). We therefore hypothesized that administering WNT sufficiently slowly would attenuate the response. Indeed, the signaling response was lower in hESCs where the WNT3A was administered gradually (**Fig. S7*A***). However, given that cells are sensitive to WNT concentration over more than two orders of magnitude (see Fig. 2), it is impractical to increase to saturating doses with a sufficiently small step size. Our experiments involved ten steps of ligand increase, and, at higher final doses of WNT, the rate of ligand increase was sufficient to saturate the response (**Fig. S7*B***). In the future it might be possible to attenuate the response to higher doses by slowly administering WNT over more extended periods with automated fluidic systems, thus more precisely probing the parameter space where hESCs are sensitive to ligand derivatives.

### WNT-target-genes have various response profiles

We next asked how WNT-target gene activation dynamics might correlate with those of ß-catenin. While some genes were transiently activated in response to WNT3A (e.g., *DKK4*, *DKK1*, and *AXIN2*), others were sustained (e.g., *EOMES*, *LEF1*) (**Fig. 4*A***). Also, since the WNT-targets *NODAL* and *BRACHYURY* are additionally regulated by Nodal signaling, we used the small molecule SB431542 to decouple the WNT response from the downstream induced Nodal response. Interestingly, Nodal is an adaptive WNT target when its self-activation is inhibited (**Fig. 4*B***). It is possible that the non-adaptive transcripts are maintained by mechanisms independent of ß-catenin, or, alternatively, their transcription is sufficiently sensitive to the level of nuclear ß-catenin following adaptation. To test whether transiently induced target genes result from the dynamics of ß-catenin, we compared the dynamics of transcription in WNT3A versus CHIR99021 treated hESCs. Induction of WNT-targets with adaptive dynamics became sustained when CHIR99021 was used (**Fig. S8**), indicating that sustaining nuclear ß-catenin is sufficient to maintain the transcription of these targets.

**Figure 4.**
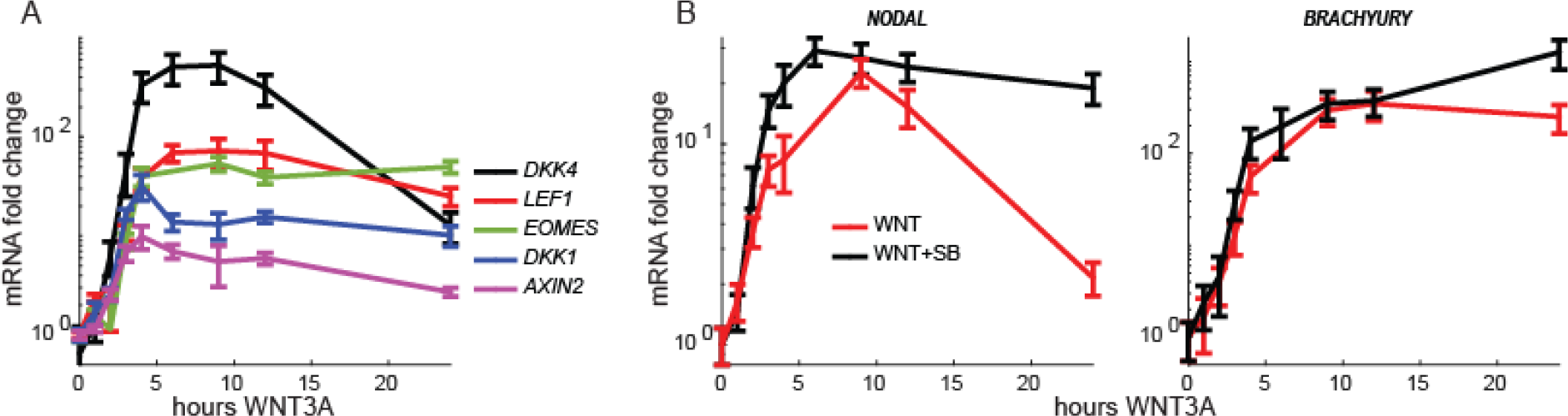
WNT Target genes have varied response profiles. (A-B) qRT-PCR in hESCs treated with (A) 300ng/ml WNT3A or (B) 300ng/ml WNT3A with or without the small molecule TGFß pathway inhibitor SB431542 (10μM) for the indicated duration.

### ß-catenin signaling dynamics are context-specific and sustained in a primitive-streak differentiation protocol

We next asked how the response to WNTs might vary across cell types. Immunofluorescent imaging using an antibody for ß-catenin revealed a variety of dynamics in response to WNT3A across different mammalian cell lines (**Fig. 5*A***), including both adaptive and sustained profiles, demonstrating that adaptation is not a universal feature of WNT-signaling and that WNT/ß-catenin dynamics can change with context and cell-fate. Since the *in vivo* equivalent of hESCs are epiblast cells, which differentiate to primitive-streak (PS) fates in response to WNT, we examined ß-catenin dynamics in hESCs differentiated to the PS-like cell-fate. For PS differentiation, we adapted a previously published protocol (18). hESCs were stimulated with Activin, BMP4, and 1 μm CHIR 99021 for 24 hours. Identity of these cells was determined by immunofluorescent staining showing downregulation of E-cadherin and SOX2 and upregulation of the PS-marker Brachyury, which confirmed the PS fate of these cells (**Fig. S9**). Interestingly, the differentiation protocol induced a sustained ß-catenin signaling profile (**Fig. 5*B*)**. This was surprising as the dose of CHIR99021 is not sufficient to induce detectable nuclear ß-catenin when presented in isolation (see Fig. 2*D*). The PS cells were refractory to exogenous WNT as adding WNT3A after 24 hours of differentiation gave no response over the mock-treated control (**Fig. 5*C***).

**Figure 5.**
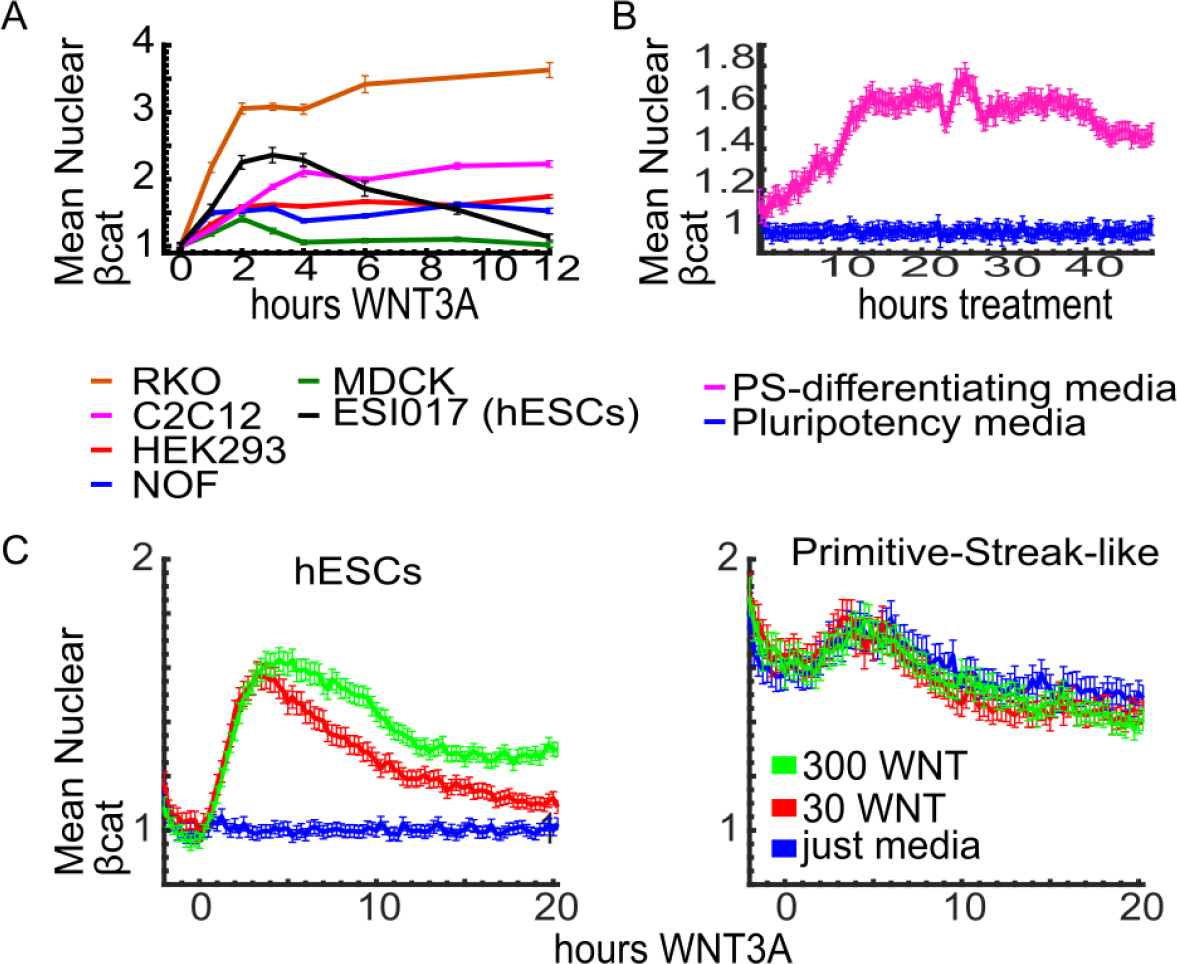
Signaling is sustained during differentiation to primitive-streak fates, and the dynamic response to exogenous WNTs varies in other cell types. (A) Quantification of anti-β-catenin by immunofluorescent imaging in the cells lines indicated in the legend. (B-C) Quantification of time-lapse movies of GFP-ß-catenin containing hESCs. (B) Treated with or without primitive-streak differentiation media. (C) Response to exogenous WNT3A in pluripotent hESCs (left) vs cells differentiated for 24 hours to primitive-streak-like fate (right). Error bars in all graphs indicate SEM of ≥ 267 cells.

### Activin and BMP activate WNT-signaling with a delay

Given the sustained induction of nuclear ß-catenin in the PS-differentiation protocol, we sought to identify how Activin and BMP might individually contribute towards ß-catenin signaling. We found that both Activin and BMP alone are capable of inducing sustained ß-catenin signaling (**Fig. 6*A*** and ***B***). ß-catenin activation with Activin and BMP was delayed compared to ß-catenin’s activation with WNT3A (**Fig. 2*B***) and SMAD activation with Activin or BMP (15), implying that ß-catenin signaling is a downstream effect of the Activin and BMP signal. Since both BMP and Activin can induce the production of new WNT ligands (13), we used IWP2 or recombinant DKK1 (an inhibitor of the canonical WNT coreceptor, LRP6) to inhibit the activity of WNTs induced downstream of Activin or BMP. We found that IWP2 completly blocked the increase in nuclear ß-catenin in response to Activin, indicating that Activin requires the presence of endogenous WNTs to activate ß-catenin signaling. However, even a combination of IWP2 and DKK1 only partially blocked the increase in nuclear ß-catenin in response to BMP, suggesting, surprisingly, that while some of the ß-catenin response to BMP is attributible to new synthesis and secretion of WNT ligands, part of the response is WNT-ligand independent.

**Figure 6.**
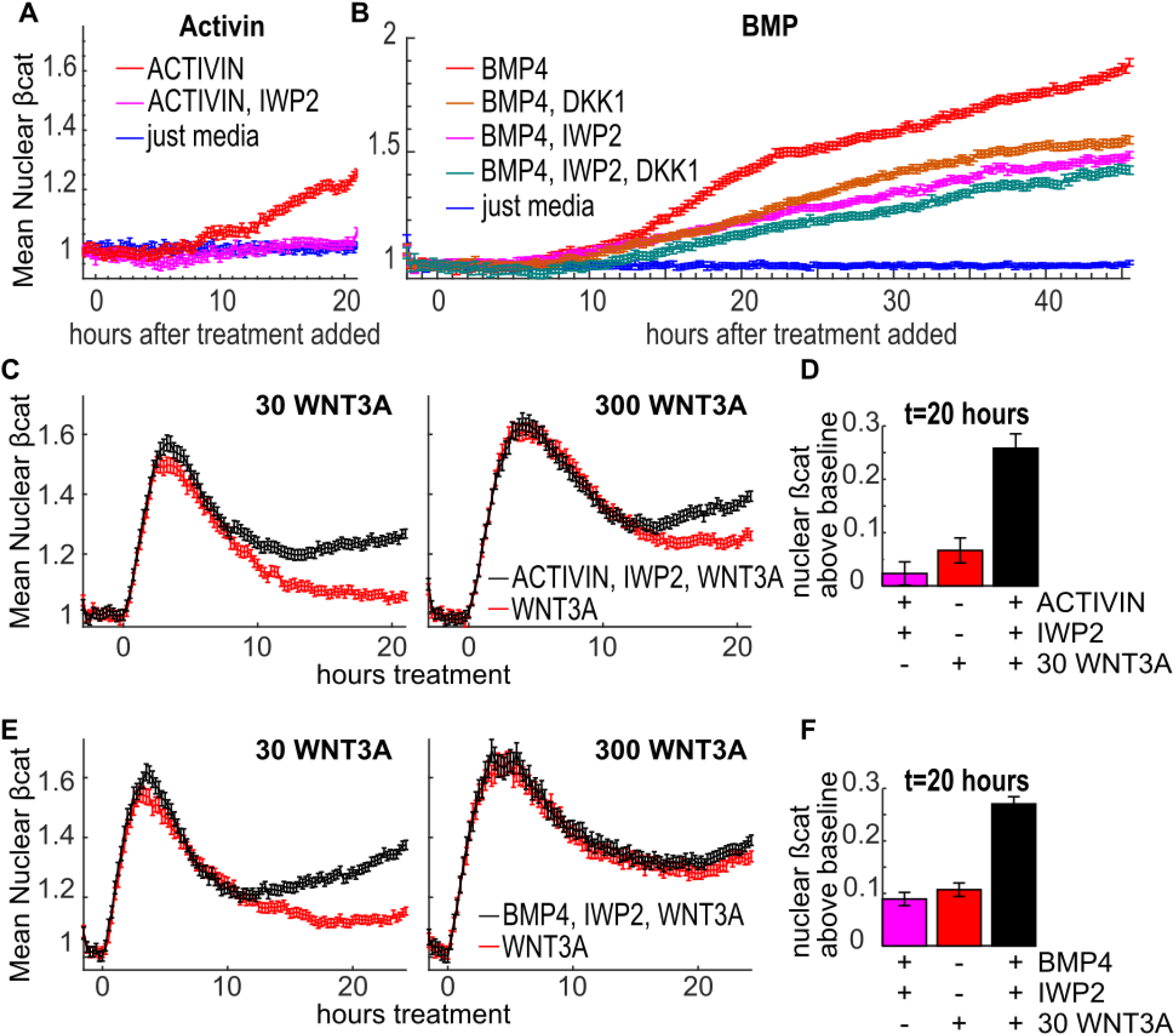
TGFß-ligands ACTIVIN and BMP4 increase ß-catenin signaling independent of WNT-ligand induction. IWP2 (3μM) is a small molecule inhibitor of endogenous WNT secretion and DKK1 (200ng/ml) is a protein that blocks canonical WNT signaling at LRP5/6 receptor. (A,B,C,E) Quantification of time-lapse imaging of GFP-ß-catenin labeled hESCs treated as indicated. All treatments are administered simultaneously. (D,F) Nuclear ß-catenin over baseline after 20 hours from indicated treatments showing TGFß-ligand synergy with WNT independent of WNT-secretion. Data is reproduced from A, C, and E. In all graphs, Activin A and BMP4 doses were 30ng/ml and 10ng/ml, respectively, and error bars represent SEM of ≥ 511 cells.

### Activin and BMP synergize with WNT, without a requirement of WNT-ligand induction

We next asked how the presence of Activin or BMP might affect the response to exogenous WNT signals. When intermediate WNT3A (30ng/ml) is added to cells at the same time as Activin (**Fig. 6*C***) or BMP4 (**Fig. 6*E***) along with IWP2, the initial response is similar to the WNT3A response alone. After 7 hours signaling continues to adapt in WNT3A only treated cells, but begins to increase in WNT3A+Activin and WNT3A+BMP treated cells. The effect of Activin or BMP4 is only partially dependent on induction of endogenous WNTs, since signaling increases despite the presence of IWP2. The total signaling after 20 hours of Activin or BMP co-treatment with WNT was more than the sum of their individual effects (**Fig. 6*D*, *F***). Activin and BMP no longer affect ß-catenin when WNT3A saturates the response (300ng/ml WNT3A, **Fig. 6*C*** and **E**). Additionally, there was little difference when ACTIVIN or BMP was added 10 hours prior to exogenous WNT3A rather than simultaneously (**Fig. S10**). This demonstrates that neither Activin nor BMP require induction of endogenous WNTs to increase ß-catenin signaling. However, while Activin still requires synergy with another source of WNTs to increase ß-catenin signaling, BMP increases ß-catenin signaling even when all other sources of WNTs have been removed.

## Discussion

Here we show that ß-catenin signaling responds adaptively to constant WNT signaling in pluripotent stem cells, and that these dynamics change dramatically with cell-context. Signaling adaptation is controlled upstream of GSK3ß by both intra- and extra-cellular mechanisms. At sub-saturating doses of WNT, extra-cellular mechanisms facilitate complete adaptation to the WNT signal, while at saturating WNT adaptation is partial and cell-autonomous. This is in contrast to TGFß signaling, which we previously showed to be adaptive in pluripotent cells (15), where adaptation is near complete even at saturating doses. Finally, our results show that TGFß and BMP signaling increase ß-catenin signaling independently of their ability to induce WNT ligands.

Our GFP-ß-catenin knock-in labeling strategy is quantitative, contains high temporal resolution, conserves spatial information, and in contrast to ß-catenin-fluorescent-protein over-expression reporters (39), maintains pathway stoichiometry. In the future we will be able to combine this approach with self-patterning differentiation assays (e.g., organoids, gastruloids, and embryoids) (40, 41), providing us with the signaling trajectories required for each cell-fate. This approach is not limited to development and can likely be used just as effectively in dissecting signaling in models of diseased tissue.

Previously, Goentoro *et al* found that total ß-catenin levels were stably increased in a rectal carcinoma cell line (RKO) which contained a mutation in E-cadherin that prevents ß-catenin localization to the membrane (42). This mutation facilitates biochemical analysis, however, it is unclear whether these dynamics are representative of other cell types. More recently, Kafri *et al* measured ß-catenin dynamics in HEK293 cells by overexpressing a YFP-ß-catenin reporter (39). While the ß-catenin dynamics we measured for RKO and HEK293 cells were within the range of dynamics measured in both of these studies, we also found that the range of dynamics was highly variable across different contexts. Additionally, RKO cells were an outlier in that of all the cell-lines tested, they exhibited the greatest fold-change in nuclear ß-catenin pre- to post-WNT treatment (see Fig. 5). Our results highlight the importance of studying ß-catenin dynamics specifically in the context of interest and caution against extrapolating between cell lines, organisms, or different differentiation states.

It is becoming increasingly appreciated that signaling pathways respond to stimulation with complex dynamics of signal transduction that influence how ligands are interpreted. For example, in hESCs, the SMAD4 response to BMP4 ligand is stable (15, 23, 43), while its response to Nodal is adaptive (15, 43). Adaptive signaling has also been observed in other pathways including EGF (44), NF-kB (45, 46), and SHH (37). Pioneering work on ERK signaling in PC12 cells have shown that ERK signals can be either transient or sustained depending on whether they are stimulated with EGF or NGF, but it remains unclear whether pathway dynamics are typically intrinsic to a particular ligand and pathway or vary depending on the context. This study demonstrates that WNT/ß-catenin signaling dynamics vary dramatically even in response to the same ligand, as we observed a range of adaptive and sustained responses depending on the stage of differentiation or the cell type. It will be interesting to revisit the dynamics of the pathways described above to determine whether context dependence is a common feature of signal transduction.

Since WNT/ß-catenin signaling controls a variety of processes in different contexts, it will be interesting to see whether certain dynamic properties are associated with distinct roles. For instance, does providing positional information during gastrulation require an adaptive response to WNTs? Are WNT/ß-catenin dynamics adaptive in similar contexts, such as anterior-posterior patterning of the neural tube? Adaptive signaling dynamics sensitize cells to the derivative of ligand concentration, and we (47, 48) and others (49, 50) have suggested that, in certain contexts, more positional information can be gained by responding to ligand derivatives compared to concentration alone. In contrast, are there scenarios where adaptation would be at odds with the role of the WNT pathway? For instance, WNT adaptation appears incompatible with the requirements for WNT in maintaining homeostasis in intestinal crypts (51, 52). In these crypts, constant WNT signaling maintains the stemness of highly proliferative multipotent cells. These cells differentiate when they move away from the WNT signal. Would adaptation not lead to the loss of stem-cells in the crypt? If cells had adapted to the WNT signal, they would be unable to determine when the signal was lost. It seems that the dynamic requirements for maintaining a constant zone might be very different than the requirements for patterning during gastrulation, and it will be interesting to compare WNT/ß-catenin dynamics between these contexts in the future.

This work revealed that additional ß-catenin signaling can be provided from TGFß and BMP through a variety of mechanisms including induction of new WNT ligands, sensitizing cells to exposure to WNT ligands, and, for BMP, a mechanism that is completely WNT ligand independent. During gastrulation, BMP signaling from the trophectoderm induces WNT signaling in the primitive streak. The WNT signal induces the TGFß ligand Nodal, which in turn activates BMP signaling. In this way, these signals are thought to reinforce each other and pattern the gastrula (13). It will be interesting to see if Activin/BMP’s induction of ß-catenin signaling outside of ligand induction is important in the context of gastrulation. One hypothesis is that TGFß or BMP mediated stabilization of WNT dynamics might be required to drive certain primitive-streak fates, and so differentiation to these fates would only occur in regions of the embryo where both signals are present. If true, it would be interesting to determine whether this synergy between pathways is independent of the induction of new WNT ligands by Nodal or BMP.

A more fine-grained investigation of the mechanisms that control WNT adaptation and WNT/TGFß/BMP pathway cross-talk is required. Are intra-cellular adaptation mechanisms always active, or are they only induced at high doses of WNT? Which proteins are involved in regulating adaptation, and are their levels modulated to control the degree of activation? More generally, future investigation is needed to understand how the dynamics of the pathway are tuned to achieve divergent functions depending on the context.

## Methods

### Cell culture, treatments, and differentiation

Human embryonic stem cells (ESIBIO; ESI017) and induced pluripotent stem cells (Coriell Institute; AICS-0058-067) were maintained in pluripotency maintenance culture as described in (23).

For all experiments with hESCs and iPSCs were seeded into mTeSR1 media (StemCell Technologies) containing rock-inhibitor Y27672 (10μM; StemCell Technologies #05875) at a density of 5×10^4^ cells/cm^2^ (except where noted otherwise). Media was changed the following morning approximately 2 hours prior to administering any treatments. Experiments with other cell lines were similar, except for seeding density and culture media. Seeding density was lowered by up to a factor of five for the largest cells, such as RKO and C2C12. DMEM (Corning #10-017) with 10% FBS (Fischer Scientific #16000044) culture media was used for C2C12, RKO, HEK293, and MDCK cell-lines. NOF151-hTERT cells were grown in a 1:1 mixture of MCDB 105 (Sigma-Aldrich #M6395) and 199 medium (Sigma-Aldrich #M0393) supplemented with 10%FBS, and 1 ng/mL epidermal growth factor (R & D Systems #PRD236)

When exogenous ligands or small molecules were administered to cells, they were pre-diluted in a volume of media that is 20% of final culture volume to facilitate rapid mixing. Recombinant proteins and small molecules used were: Activin A (R & D Systems 338-AC; 30ng/ml), BMP4 (R & D Systems #314BP050; 10ng/ml), CHIR99021 (MedChem Express #HY-10182), DKK1 (R & D Systems #5439-DK-010; 300ng/ml), IWP2 (Stemgent #04-0034; 3μM), SB431542 (StemCell Technologies #72232; 10μM), WNT3A (R & D Systems #5036-WN), WNT3A packaged with WNT-stabilizer reagent (AMSBIO #AMS.rhW3aL-002-stab), Y-27632 (StemCell Technologies #72302; 10μM)

For WNT ramps in supplemental figure 8, WNT3A was administered every two hours so that the total administered WNT after each step was as indicated. Ramp duration and time between steps was chosen based on experimental feasibility.

PS differentiation conditions in figure 5 are adapted from (53). Our modifications were using Essential 6 Media (Gibco #A15165-01) as a base and lowering CHIR99021 dose to 1μM. The other supplements were as described by Loh *et al*: 30ng/ml Activin A, 20ng/ml FGF2 (Life Technologies; PHG6015), and 40ng/ml BMP4.

### Plasmids and Generation of GFP-β-catenin labeled hESC cell line

We followed the protocol in (19) for CRISPR-cas gene editing. Guide RNA expression was from a PCR amplified gBlock from IDT as described in (19) but containing the ß-catenin targeting sequence, cgtggacaatggctactca, located close to the ATG start codon. The homology donor template DNA was prepared by PCR amplifying PuroR-T2A-GFP over two rounds. The primers in the first round contained 5’ overhangs which add part of the ß-catenin homology arms. The second round of PCR is done using the PCR product of the first reaction as a template, and uses primers containing the remaining homology arm sequence in their 5’ overhangs. Cas9 expression plasmid, homology donor DNA, and guide RNA were co-nucleofected in hESCs using P3 Primary Cell 4D-Nucleofector^®^ X Kit (Lonza #V4XP-3012) and positive transformants were selected with puromycin (2ug/ml; Life Technologies # A1113803). RFP-H2B construct was described in (23). GFP and RFP double positive cells were obtained with FAC-sorting.

#### Sequence of gBlock for ß-catenin guideRNA

AGTATTACGGCATGTGAGGGCCTATTTCCCATGATTCCTTCATATTTGCATATACGATACAAGGCTGTTAGAGAGATAATTGGAATTAATTTGACTGTAAACACAAAGATATTAGTACAAAATACGTGACGTAGAAAGTAATAATTTCTTGGGTAGTTTGCAGTTTTAAAATTATGTTTTAAAATGGACTATCATATGCTTACCGTAACTTGAAAGTATTTCGATTTCTTGGCTTTATATATCTTGTGGAAAGGACGAAACACCGcgtggacaatggctactcaGTTTAAGAGCTATGCTGGAAACAGCATAGCAAGTTTAAATAAGGCTAGTCCGTTATCAACTTGAAAAAGTGGCACCGAGTCGGTGCTTTTTTGTTTTAGAGCTAGAAATAGCAAGTTAAAATAAGGCTAGTCCGTTTTTAGCGCGTGCGCCAATTCTGCAGACAAATGGCTCTAGAGGTACGGCCGCTTCGAGCAGACATGATAAGATACATTGA

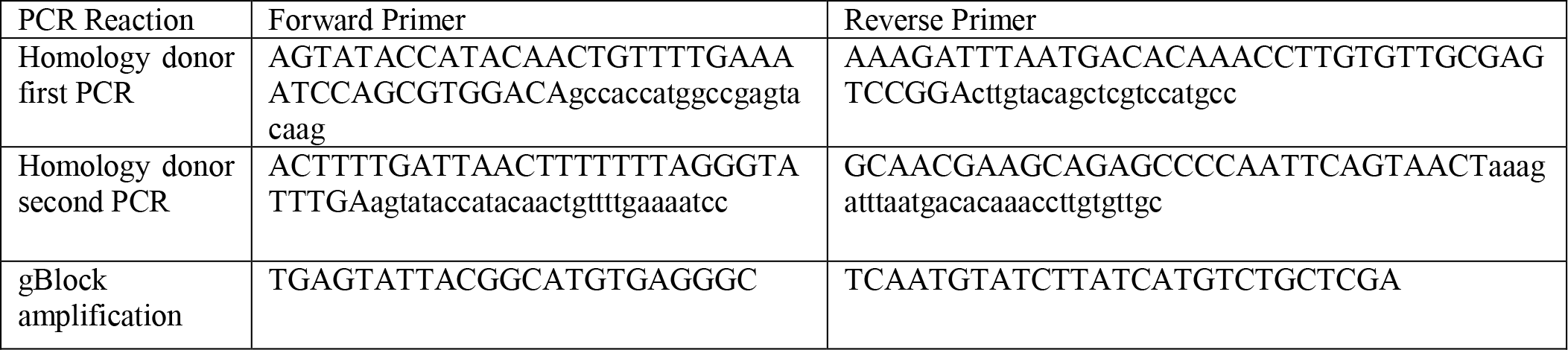

### Immuo-fluorescent antibody staining

Cells were fixed with 4% PFA and stained as in(23). Antibodies and dilutions were used as follows:

Immuo-fluorescent antibody staining Cells were fixed with 4% PFA and stained as in(23). Antibodies and dilutions were used as follows:
| Protein | Species | Dilution | Vendor | Catalog number |
| --- | --- | --- | --- | --- |
| β-catenin | Mouse | 1:200 | BD Biosciences | 610154 |
| E-cadherin | Mouse | 1:200 | Fisher Scientific | 3195S |
| Brachyury | Goat | 1:300 | Fisher Scientific | AF2085 |
| Sox2 | Rabbit | 1:200 | Fisher Scientific | 5024S |

### Western Blots

Western blots were performed using standard protocols. Antibodies used: β-catenin (mouse; BD Biosciences #610154; dilution 1:20000), GFP (rabbit; Cell Signaling #2956S; dilution 1:1000), Peroxidase-Beta-Actin (Sigma-Aldrich #A3854; 1:50000), Peroxidase-Rabbit-IgG (Jackson Immuno Research #711-035-152; 1:2500), Peroxidase-Mouse-IgG (Jackson Immuno Research #711-035-150; 1:5000)

### RT-qPCR

RT-qPCR was performed using manufacturer methods. Briefly, RNAqueous-Micro Total RNA Isolation Kit (Fisher Scientific; AM1931) was used to prepare RNA, and SuperScript Vilo cDNA Synthesis Kit (Fisher Scientific; 11754-050) was used to prepare cDNA. qPCR reactions were done on a Step One Plus Real-Time PCR System (Applied Biosystems) using Power SYBR Green PCR Master Mix (Life Technologies; 4367659). ATP5O was used to normalize all genes. Primers are below:

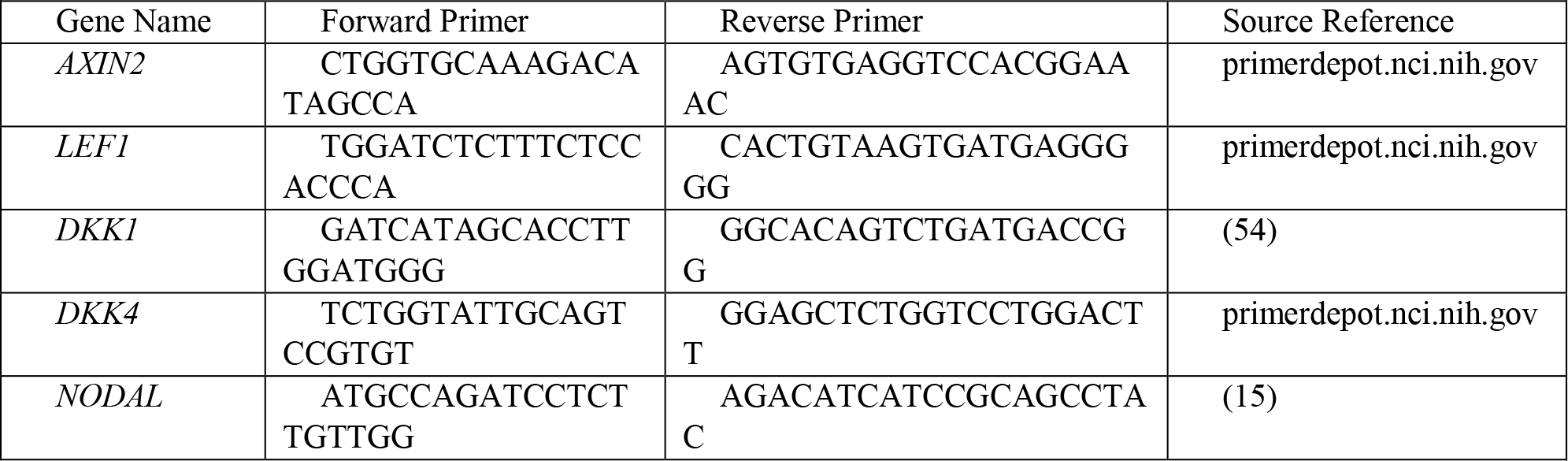

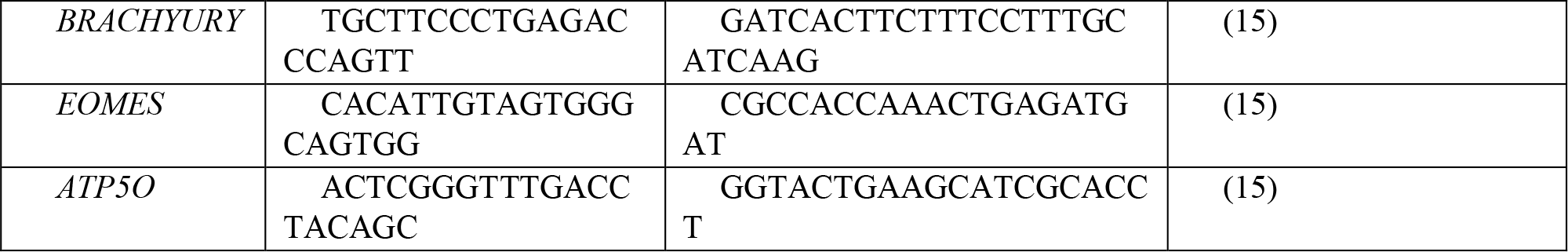

### Imaging and analysis

Imaging was performed on an Olympus/Andor spinning disk confocal microscope using either a 20X, 0.75 Na air or a 40X, 1.25 Na silicon oil. Most images displayed in manuscript were taken at 40X, while the majority of movies were quantified at 20X (see supplemental discussion). Time-lapse imaging intervals were either 10 or 15 minutes, and z-stacks were acquired in 3 planes spaced 2.5 micrometers apart. Image analysis was done combining custom software written in MATLAB (Mathworks) and described in (24) (available at www.github.com/josephkm) and Ilastik (55) (www.ilastik.org). Briefly, max intensity projections were taken across z-slices, and background was subtracted. Background was identified by minimum intensity projection across many images, and was manually checked for consistency. Nuclei pixels were identified using Ilastik, and resulting masks were imported to MATLAB for segmentation of cells and image quantification. Nuclear intensities of each cell were normalized against that cellʼs nuclear marker, and the mean and standard error of these cells was then normalized against the negative control for that experiment (typically a mock-treated control group). For live-imaging datasets, a median filter in time over a window of 12 time steps was applied to the control condition before normalization to avoid propagation of fluctuations from the control to the experimental condition. The control traced in each figure shows the control condition normalized to its own median filter so that the variability in this condition can be observed.

## Acknowledgements

This work was funded by grants from NIGMS (R01GM126122), NSF (MCB-1553228), CPRIT (RR140073), and the Simons Foundation (511079). We thank Sapna Chhabra for help with micropatterned gastruloid experiments, and members of AW lab for helpful discussions and comments on the manuscript.

